# Meiotic DNA exchanges are promoted by proximity to the synaptonemal complex

**DOI:** 10.1101/2022.08.11.503613

**Authors:** David E. Almanzar, Spencer G. Gordon, Chloe Bristow, Lexy von Diezmann, Ofer Rog

**Author notes:** equally contributing authors (Spencer G. Gordon, Chloe Bristow).

## Abstract

During meiosis, programmed double strand DNA breaks are repaired to form exchanges between the parental chromosomes called crossovers. Chromosomes lacking a crossover fail to segregate accurately into the gametes, leading to aneuploidy. Crossover formation requires the promotion of exchanges, rather than non-exchanges, as repair products. However, the mechanism underlying this meiosis-specific preference is not fully understood. Here, we study the regulation of meiotic sister chromatid exchanges in *Caenorhabditis elegans* by direct visualization. We find that a conserved chromosomal interface that promotes exchanges between the parental chromosomes, the synaptonemal complex, also promotes exchanges between the sister chromatids. In both cases, exchanges depend on recruitment of the same set of pro-exchange factors to repair sites. Surprisingly, while the synaptonemal complex usually assembles between the two DNA molecules undergoing an exchange, its activity does not rely on a specific chromosome conformation and it can also promote sister exchanges when assembling next to the sisters. This suggests that the synaptonemal complex regulates exchanges by establishing a nuclear domain conducive to nearby coalescence of exchange-promoting factors.

## Introduction

Meiosis is the specialized cell division that creates haploid gametes from diploid precursor cells. For the paternal and maternal copies of each chromosome (homologs) to segregate accurately, they must form at least one reciprocal inter-homolog exchange (crossover). At the same time, the complex DNA acrobatics that generate crossovers carry a risk of corrupting the genome. As a result, crossover formation is tightly regulated.

Crossovers form when programmed DNA double strand breaks are repaired by homologous recombination, which restores genomic integrity using information from a template that shares sequence homology. While homologous recombination throughout development uses many of the same molecular actors and DNA intermediates, their regulation during meiosis is distinct in two respects. The first is the choice of repair template: while mitotic cells use the identical copy available in the sister chromatid, meiotic cells preferentially use the similar homolog as a repair template (Humphryes & Hochwagen, 2014). The second is the nature of the repair products. Repair events can be resolved to reciprocally exchange flanking sequences or as non-exchange products that locally patch the break. Exchanges jeopardize both the broken DNA molecule and the template and are indeed avoided in mitotic cells where they are associated with cancer predisposition (Chaganti *et al*, 1974). In meiosis, however, a tightly regulated number of repair intermediates (in *C. elegans*, exactly one per chromosome) are processed as exchanges to form crossovers, with most other repair events resolved as non-exchanges (Pyatnitskaya *et al*, 2019).

The mechanism that promotes formation of a precise number of crossovers while channeling other repair events into a non-exchange fate depends on a set of conserved pro-crossover factors known as the ZMMs (named after the budding yeast proteins <u>Z</u>ip1-4, <u>M</u>er3, <u>M</u>sh4-5 and Spo16; the *C. elegans* components, also referred to here as ZMMs, are ZHP-1-4, COSA-1 and MSH-5-HIM-14^MSH-4^; (Pyatnitskaya *et al*, 2019)). These factors form foci at a subset of inter-homolog repair sites – also called recombination nodules – that correspond in their number and distribution to crossovers (Yokoo *et al*, 2012; Carpenter, 1975). Recombination nodules designate crossovers by regulating the processing of repair intermediates localized within them to yield exchanges (Pyatnitskaya *et al*, 2019).

Key to the regulation of recombination nodules is a conserved chromosomal interface - the synaptonemal complex – that localizes to between the homologs. Once the homologs independently assemble their chromatin around a backbone called the axis and find each other (“pair”), they are aligned along their length by the central region of the synaptonemal complex (SC-CR; (MacQueen *et al*, 2002; Rog & Dernburg, 2013)). Both DNA breaks and the SC-CR are necessary for formation of recombination nodules in worms: when a chromosome lacks breaks or fails to assemble SC-CR, it does not form recombination nodules and does not undergo crossovers (Rosu *et al*, 2013; Stamper *et al*, 2013; Yokoo *et al*, 2012). However, conditions that eliminate breaks or the SC-CR throughout the nucleus reveal that each contributes independently to recombination nodule formation. In animals that lack the SC-CR, ZMM proteins form break-dependent foci on chromosomes (Li *et al*, 2018; Cahoon *et al*, 2019); and in the absence of breaks, ZMM proteins form a focus abutting chromatin-free aggregates of SC-CR material (Shinohara *et al*, 2015; Rog *et al*, 2017; Zhang *et al*, 2018; Tsubouchi *et al*, 2006). However, the molecular mechanisms by which the SC-CR and DNA breaks control crossover formation remain poorly understood.

Recently, we showed that exchanges between sister chromatids are rare in *C. elegans* (Almanzar *et al*, 2021). Interestingly, we found that the SC-CR can promote sister exchanges when it mis-localizes to unpaired chromosomes (Cahoon *et al*, 2019; Almanzar *et al*, 2021). This occurred concomitantly with formation of recombination nodules, raising the possibility that recombination nodules also promote exchanges between sisters.

Here, we study the mechanism by which the SC-CR promotes sister exchanges. By testing several conditions where the SC-CR assembles on chromosomes that exclusively undergo sister-directed repair, we found elevated rates of sister exchanges that correspond to the number of recombination nodules. Recombination nodules are necessary and sufficient for high levels of sister exchanges, suggesting that SC-CR-mediated coalescence of ZMM proteins promotes all exchanges, whether they occur between homologs or sisters. Furthermore, we find that sister exchanges occur when the SC-CR assembles both between the sisters or next to them, suggesting that proximity to the SC-CR promotes formation of nearby recombination nodules.

## Results

### The SC-CR promotes sister exchanges

Previously, we used a novel approach to cytologically score sister exchanges in conditions where the homolog was unavailable and therefore the sister was used as the template for homologous recombination. We found that, while sister exchanges were rare in *him-8* worms where the X chromosome does not pair or associate with the SC-CR, they were common in three scenarios where the SC-CR associates with unpaired chromosomes: deletion of the cohesin subunit *rec-8*, and mutations in SC-CR subunits *syp*□*3(me42)* and *syp*□*1*^*K42E*^ (Almanzar *et al*, 2021).

To rule out an X chromosome-specific effect on sister exchanges, we examined *zim-2* animals, where chromosome V does not pair or associate with the SC-CR, in an analogous fashion to the X chromosome in *him-8* mutants (Phillips & Dernburg, 2006). We found sister exchanges on the unpaired chromosomes to be a relatively rare outcome of DNA repair in *zim-2* animals (Fig. 1), albeit slightly elevated compared with *him-8* animals: 10.0% versus 4.2% sister exchanges per unpaired chromosome (Pearson’s chi-square, p = 0.00351). The elevated level of sister exchanges in *zim-2* worms is likely caused by a nucleus-wide increase in the number of breaks that are induced in response to unpaired chromosomes, rather than a chromosome-specific effect on repair. This idea is consistent with elevated sister exchanges on both paired and unpaired chromosomes in *zim-2* animals (see Fig. 4, below), and with the more numerous foci of the DNA repair protein RAD-51 – a proxy for the number of breaks – in *zim-2* versus *him-8* animals (7.87 versus 5.49 foci, p <0.0001, Student’s t-test; Fig. S1; (Colaiácovo *et al*, 2003)).

**Figure 1:**
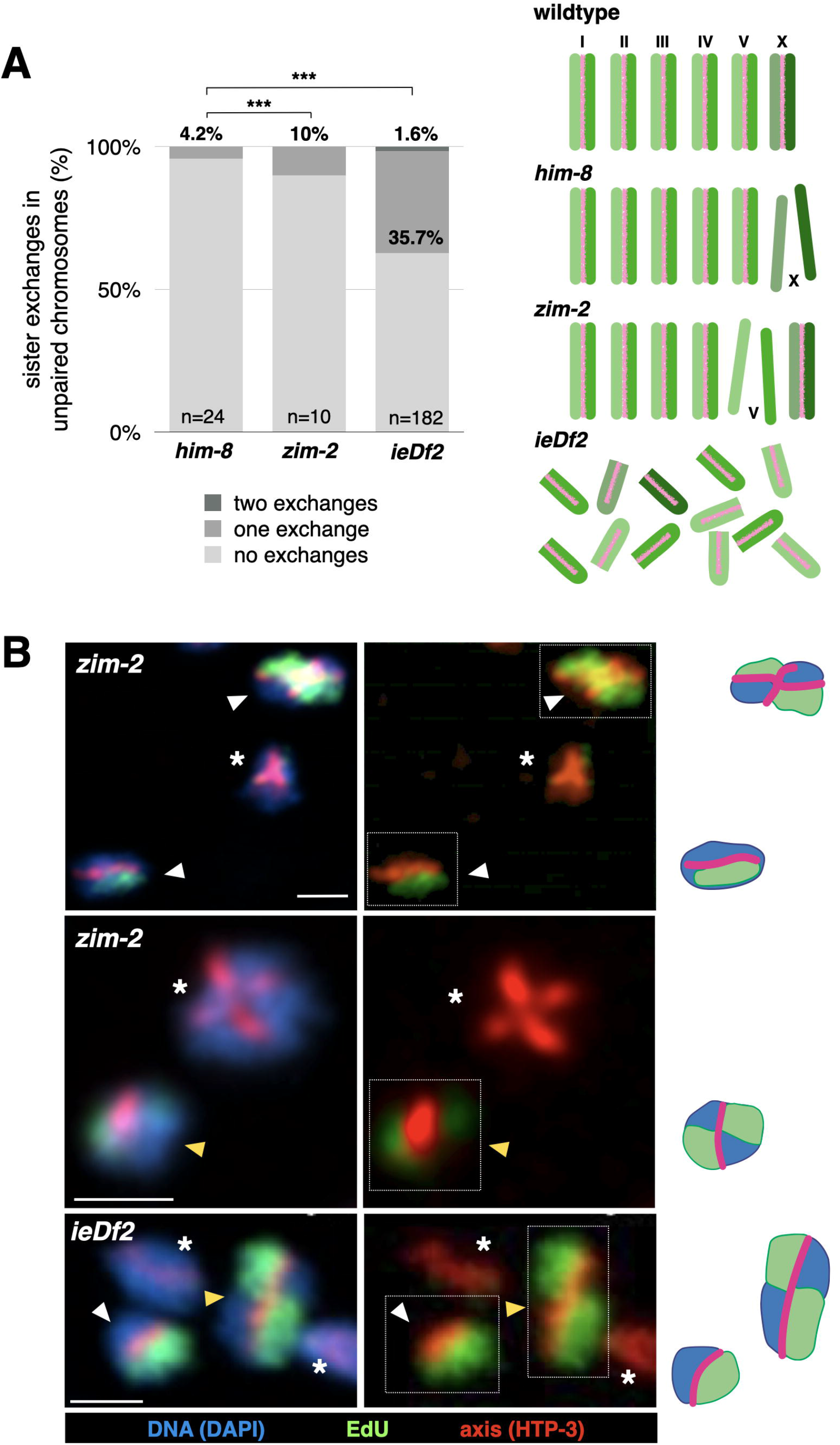
The SC-CR promotes sister exchanges when it localizes to unpaired chromosomes. A. Exchanges are elevated on unpaired chromosomes in *ieDf2* animals, but not in *him-8* and *zim-2* animals. Pairwise comparisons between *him-8* and *zim-2* were significant (p = 0.00351, Pearson’s chi-square). Pairwise comparisons between *him-8, ieDf2* and *syp-1*^K42E^ animals were all significant (p < 0.0001, Pearson’s chi-square). *him-8* data is taken from (Almanzar *et al*, 2021). Diagrams of the different genotypes are shown to the right. Chromosomes are shown in green and the SC-CR in pink. Note that the unpaired chromosomes in *him-8* and *zim-2* animals (the X chromosome and chromosome V, respectively), are not associated with the SC-CR, whereas all chromosomes are unpaired and associated with the SC-CR in *ieDf2* animals. B. Representative images and interpretive diagrams of exchange and non-exchange chromosomes in *ieDf2* and *zim-2* animals. Yellow arrows denote exchange chromatids, white arrows denote non-exchange chromatids, and asterisks denote unlabeled chromatids (not scored). Interpretive diagrams of chromosomes surrounded by dashed white boxes are shown to the right. Red, axis (anti-HTP-3 antibodies); green, EdU; blue, DNA (DAPI). Note the EdU signal crossing the axis in exchange chromatids. Scale bars = 1µm.

The conditions we examined previously – *rec-8, syp-3(me42)* and *syp-1*^*K42E*^ animals – mis-localize a mutated synaptonemal complex. To test whether a synaptonemal complex made of unaltered components can also promote sister exchanges we examined *ieDf2* worms, where all chromosomes fail to pair. In contrast to mutants like *him-8* or *zim-2*, unpaired chromosomes in *ieDf2* worms undergo ‘fold-back’ that brings together the left and right arms of each chromosome with the SC-CR assembling between them (Harper *et al*, 2011). We observed 37.3% sister exchanges per chromosome in *ieDf2* worms (Fig. 1), significantly higher than in *him-8* animals (p < 0.000005, Pearson’s chi square).

### Recombination nodules promote sister exchanges

The formation of crossovers is accompanied by the coalescence of ZMM proteins to recombination nodules (Pyatnitskaya *et al*, 2019), leading us to hypothesize that designation of an exchange fate for inter-sister repair events occurs in recombination nodules on unpaired chromosomes.

We first confirmed that recombination nodules, marked by the tagged ZMM protein GFP-COSAL1, are associated with sister exchanges in the above-mentioned conditions. In *ieDf2* animals we observed an average of 4.7 recombination nodules per nucleus, similar to the number of sister exchanges: 4.6 (Fig. 2A-B). In *syp-3(me42)* animals, there were 7.1 sister exchanges and 7.4 recombination nodules per nucleus, and in *syp-1*^K42E^ animals there were 10.1 sister exchanges corresponding to an average of 16.4 recombination nodules per nucleus (Fig. 2A-B; (Gordon *et al*, 2021; Almanzar *et al*, 2021); see Discussion for potential explanation for the latter discrepancy). In *rec-8* animals, an average of 6.8 sister exchanges were accompanied by 10.4 recombination nodules per nucleus (Almanzar *et al*, 2021; Cahoon *et al*, 2019). In *him-8* animals, consistent with the rare sister exchanges on the unpaired X chromosomes – only 4.2% underwent an exchange – recombination nodules were absent on the X chromosome (p < 0.00001, Student’s t-test), and we observed an average of 5.2 recombination nodules per nucleus, one on each of the five autosomes.

**Figure 2:**
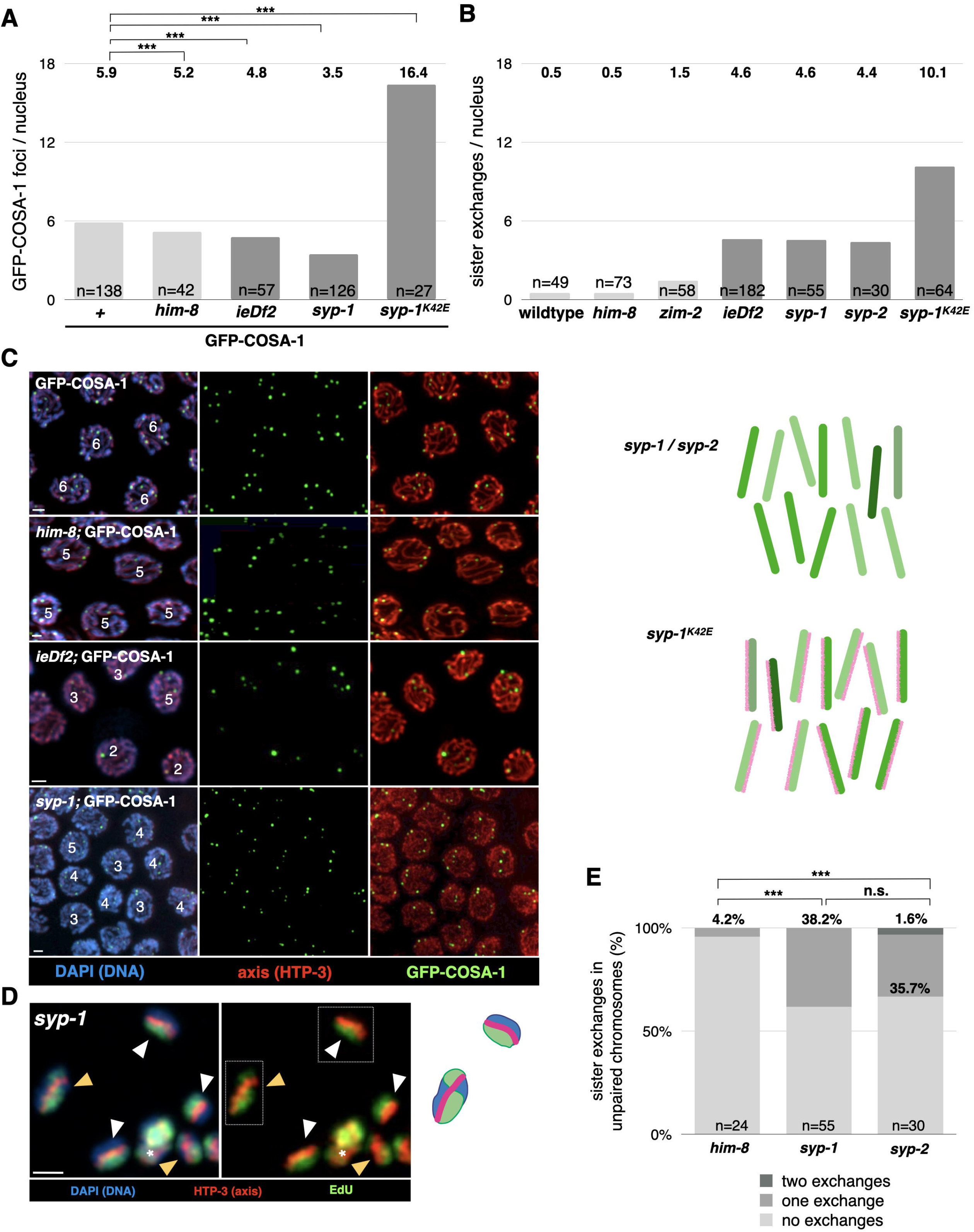
Sister exchanges correspond with recombination nodules. A. Average number of GFP-COSA-1 foci in late pachytene nuclei of animals of the designated genotype. Results were significantly different from wildtype for all genotypes (p<0.000001, Student’s t-test). n value indicates the number of nuclei counted. Data for *syp-1*^K42E^ is from (Gordon *et al*, 2021). Light gray shading indicates conditions where only one chromosome is unpaired; dark gray shading indicates all chromosomes are unpaired. B. Total number of sister exchanges extrapolated from the exchan 4A and (Almanzar *et al*, 2021). Shading is as in panel A. Note correspondence between the values in panels A and B for conditions where all chromosomes are unpaired. C. Representative images of late pachytene nuclei in wildtype, *him-8, ieDf2* and *syp-1* animals that also carry the GFP-COSA-1 transgene. Red, axis (anti-HTP-3 antibodies); green, GFP-COSA-1 (anti-GFP antibodies); blue, DNA (DAPI). Scale bar = 1µm. Diagrams of the different genotypes are shown to the right. Chromosomes are shown in green and the SC-CR in pink. Note that while the all chromosomes are unpaired in both scenarios, they are associated with the SC-CR only in *syp-1*^K42E^ animals. D. Representative images of exchange and non-exchange chromatids in *syp-1* animals. Yellow arrows denote exchange chromatids, white arrows denote non-exchange chromatids, and asterisks denote unlabeled chromatids (not scored). Interpretive diagrams of chromosomes surrounded by dashed white boxes are shown to the right. Red, axis (anti-HTP-3 antibodies); green, EdU; blue, DNA (DAPI). Scale bar = 1µm. E. Sister exchanges are elevated upon removal of the SC-CR. Pairwise comparisons between *him-8* and *syp-1* or *syp-2* animals were significant (p=0.0046 and p=0.0212, respectively, Pearson’s chi-square test).

To test whether recombination nodules are sufficient for promoting sister exchanges, we analyzed worms lacking an SC-CR, where ZMMs are ectopically recruited to DNA breaks on unpaired chromosomes. In the absence of SC-CR, an average of 3.5 recombination nodules formed on unpaired chromosomes (Fig. 2B; as was observed in (Cahoon *et al*, 2019; Li *et al*, 2018)), similar to the number of sister exchanges: an average of 4.6 sister exchanges in *syp-1* and 4.0 in *syp-2* (Fig. 2C-E). These numbers are significantly higher than those observed in *him-8*, where recombination nodules do not form on the unpaired chromosomes (p-value < 0.00001 in both pairwise comparisons, Pearson’s chi-square).

### Most sister exchanges depend on recombination nodules

To test whether sister exchanges require recombination nodules, we used *syp-1*^K42E^ animals grown at 25°C, where sister exchanges are common. We conditionally depleted the ZMM factor ZHP-3, which is essential for crossovers, by growing *zhp-3-FLAG-AID* in the presence of auxin (hereafter *zhp-3(-)*; (Zhang *et al*, 2015, 2018; Jantsch *et al*, 2004; Bhalla *et al*, 2008)). As expected, *syp-1*^K42E^ *zhp-3(-)* animals lacked recombination nodules (Fig. 3A), and, compared with *syp-1*^K42E^ animals, underwent significantly fewer sister exchanges: 3.8 *versus* 10.1 exchanges per nucleus in *syp-1*^K42E^ *zhp-3(-)* and *syp-1*^K42E^, respectively (p = 0.00007, Pearson’s chi-square; Fig. 3B-C). The effect of ZMM depletion suggests that most sister exchanges occur in recombination nodules.

**Figure 3:**
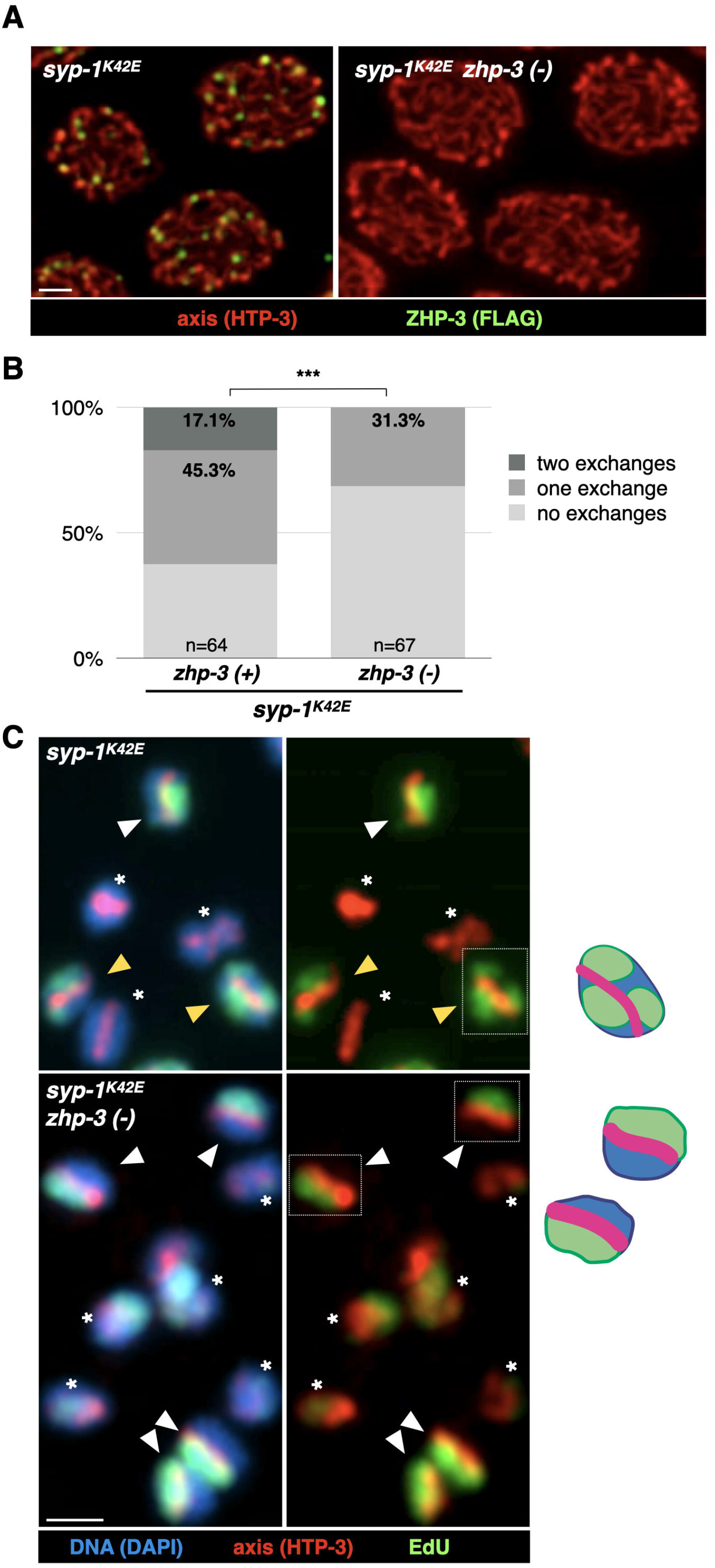
Most sister exchanges in *syp-1*^K42E^ animals depend on ZHP-3. A. Representative pachytene nuclei in *syp-1*^K42E^ *zhp-3-FLAG-AID* with and without auxin (left and right, respectively). Note the absence of recombination nodules – foci of ZHP-3 – when grown on auxin. Red, axis (anti-HTP-3 antibodies); green, ZHP-3 (anti-FLAG antibodies); blue, DNA (DAPI). Scale bars = 1µm. B. Most sister exchanges in the syp*-1*^K42E^ background are dependent on ZHP-3. Sister exchanges in *syp-1*^K42E^ animals grown with or without auxin are significantly different (p<0.0001, Pearson’s chi-square test). Data for *zhp-3(+)* animals includes data from (Almanzar *et al*, 2021). C. Representative images of exchange and non-exchange chromosomes in *syp-1*^K42E^. Yellow arrows denote exchange chromatids, white arrows denote non-exchange chromatids, and asterisks denote unlabeled chromatids (not scored). Interpretive diagrams of chromosomes surrounded by dashed white boxes are shown to the right. Red, axis (anti-HTP-3 antibodies); green, EdU; blue, DNA (DAPI). Scale bars = 1µm.

The factors responsible for the remaining sister exchanges in *syp-1*^K42E^ *zhp-3(-)* worms are unknown. ZMM-independent, or ‘class II’, crossovers have been observed in other species (Gray & Cohen, 2016), but their existence and functional relevance in *C. elegans* is a matter of debate (Youds *et al*, 2010). Our finding that sister exchanges can form without recombination nodules in certain mutant scenarios suggests that these exchanges may be generated by the same mechanisms responsible for class II crossovers.

### Sister exchanges and crossovers are regulated together

One of the most intriguing features of meiotic crossover regulation is the inhibitory effect a crossover exerts on nearby inter-homolog repair intermediates on the same chromosome (von Diezmann & Rog, 2021; Muller, 1916; Sturtevant, 1913). Termed crossover interference, this regulation is particularly robust in *C. elegans*, where it acts over micro-scale distances to yield exactly one recombination nodule, and therefore crossover, per chromosome (Libuda *et al*, 2013). Since recombination nodules are required for both crossovers and sister exchanges, we wondered whether they interfere with one another.

To test this hypothesis, we analyzed animals that are heterozygous for *nT1*, a large reciprocal translocation between chromosomes IV and V. In *nT1/+* animals, chromosomes IV and V each assemble SC-CR along their entire length but are homologously paired along only 15% of chromosome IV and 39% of chromosome V (MacQueen *et al*, 2005). The rest of chromosomes IV and V are juxtaposed with non-homologous sequences, so only sister-directed repair is possible. We found that the crossovers that form on the homologously-paired portions suppress sister exchanges and formation of additional recombination nodules along the entire chromosome: sister exchanges were not elevated in this background (Fig. 4A-B; compared to *him*□*8*, p = 0.796, Pearson’s chi-square), and we observed no more than one recombination nodule per chromosome (an average of 5.8 per nucleus; Fig. 4C-D). This indicates that crossovers exert inhibitory effects over sister exchanges *in cis* along the entire length of the chromosome, in a similar manner to the inhibitory effect exerted over additional crossovers on the same chromosome.

**Figure 4:**
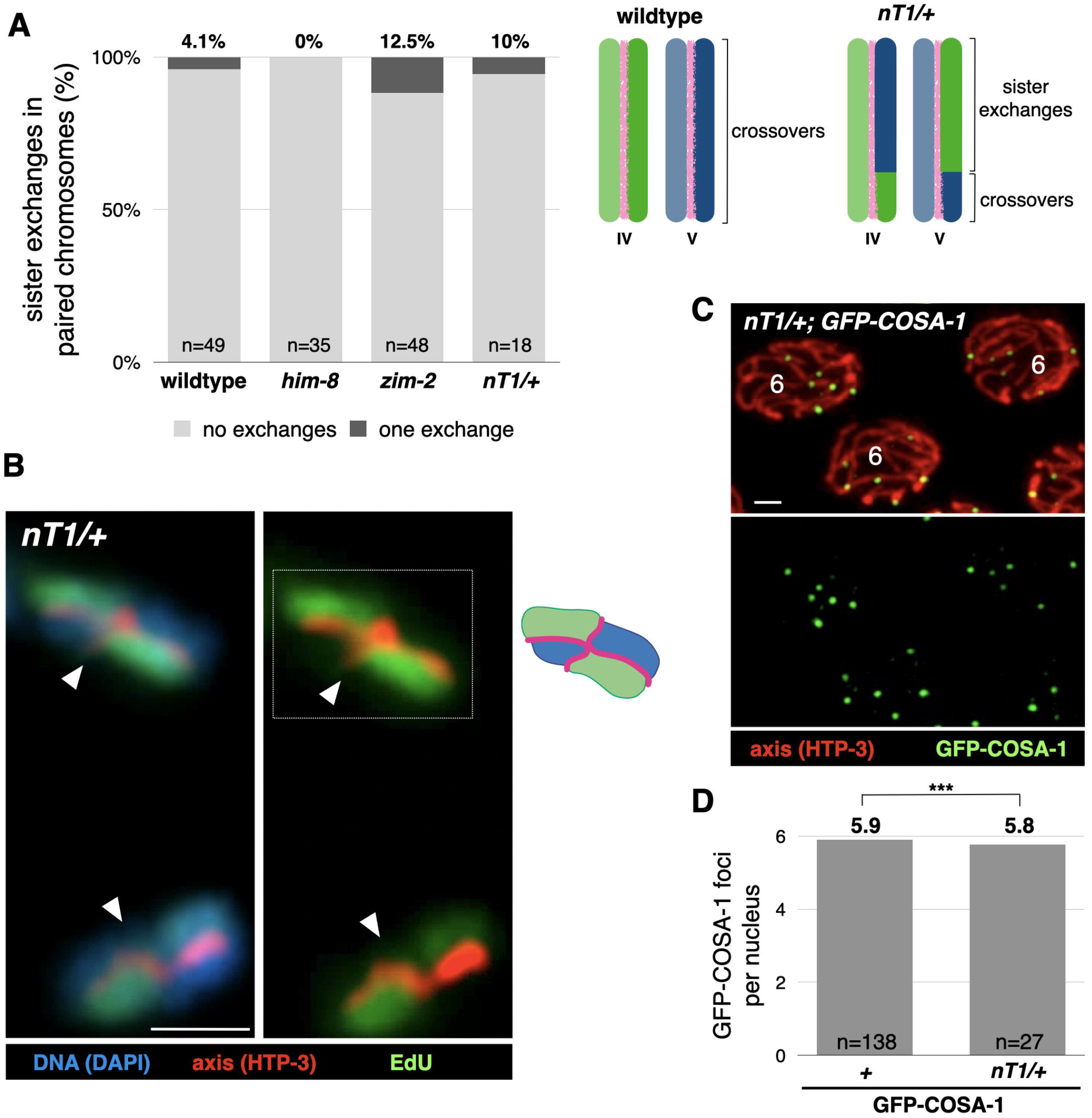
Sister exchanges and crossovers are regulated together. A. Sister exchanges remain rare when part of the chromosome can only undergo sister exchanges. Pairwise comparisons of exchanges on paired chromosomes between wildtype, *him-8, zim-2* and *nT1/+* animals were not significant (P > 0.05 in all examples, Pearson’s chi-square). Diagrams of chromosomes IV and V in wildtype and *nT1/+* animals are shown in blue and green, respectively, with the SC-CR in pink. In *nT1/+* animals, crossovers are only possible in the homologously-paired bottom portions of the chromosomes. B. Representative partial projections of exchange and non-exchange chromosomes in *nT1/+* animals and interpretive cartoons. Asterisks denote unlabeled or unscored chromatids. Red, axis (anti-HTP-3 antibodies); green, EdU; blue, DNA (DAPI). Scale bar = 1µm. C. Representative images of late pachytene nuclei in *nT1/+* animals that also carry the GFP-COSA-1 transgene. Interpretive diagrams of chromosomes surrounded by dashed white boxes are shown to the right. Red, axis (anti-HTP-3 antibodies); green, GFP-COSA-1 (anti-GFP antibodies); blue, DNA (DAPI). Scale bar = 1µm. D. Quantification of GFP-COSA-1 foci in wildtype and *nT1/+* animals. Results were significant between genotypes (p=0.0025, Student’s t-test), although the magnitude of the effect suggests it is not biologically significant. n value indicates the number of nuclei counted.

### The SC-CR promotes sister exchanges in various conformations

During crossover formation, the SC-CR assembles between the two homologs, and consequently, between the two DNA molecules being exchanged. The near-universal conservation of their relative positions suggests that the SC-CR may promote exchanges by placing the swapped DNA molecules across from each other. However, it is challenging to test this idea since the SC-CR plays multiple roles in crossover formation, including bringing the homologs together. Our finding that the SC-CR can promote all meiotic exchanges allows to address this possibility by examining exchanges between the sisters, which are held together independently of the SC-CR.

We used STED super-resolution microscopy to probe the conformation of the SC-CR relative to axes, where chromatin is tethered. We first examined wildtype animals, where the enhanced resolution confirmed that the SC-CR places the axes of the two homologs parallel to one another, separated by 160 nm (Fig. 5; (Goldstein & Slaton, 1982; Köhler *et al*, 2017; Rog & Dernburg, 2013)).

**Figure 5:**
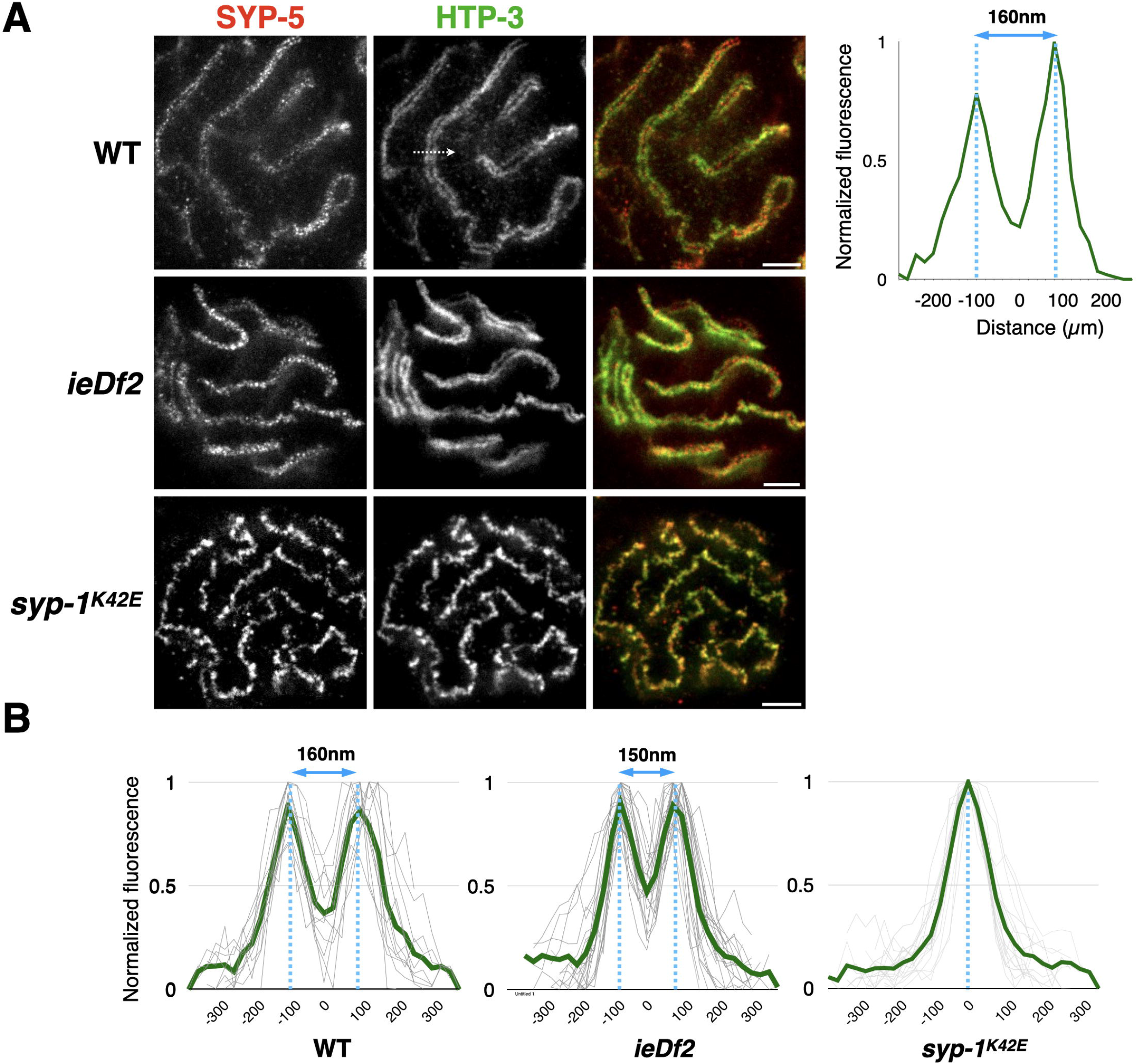
SC-CR in various conformations can promote sister exchanges. A. Representative STED images of pachytene nuclei from wildtype, *ieDf2* and *syp-1*^K42E^ worms. Red, SYPL5; green, HTPL3. Dashed white arrow denotes line scan shown to the right. HTP-3 fluorescence is normalized such that maximum fluorescence is 1. Note that the SC-CR localizes between the axis in both wildtype and *ieDf2* worms, but the SC-CR and axis signals colocalize in *syp-1*^K42E^ worms. See Figs. 1 and 2 for diagrams of the different genotypes. Scale bars = 1µm. B. Overlayed line scans show similar distribution in wildtype (n = 12) and *ieDf2* (n = 27) worms – two axis peaks separated by 150-160nm – while *syp-1*^K42E^ worms (n = 18) show a single peak. Averages are shown in green.

We next analyzed conditions where the homologs are not paired, and the SC-CR instead promotes sister exchanges. Analysis of *ieDf2* worms revealed a conformation that resembles the wildtype, with the two axes separated by 150 nm and the space between them occupied by the SC-CR (Fig. 5). Crucially, however, since the two axes in this case belong to the left or right arms of the chromosome, the two sisters being exchanged are situated to one side of the SC-CR. In *syp*□*1*^*K42E*^ animals, however, we observed the SC-CR and the axis – where the two sisters are tethered – co-localizing to form a single overlapping thread (Fig. 5). In this case the two exchanged DNA molecules are localized outside the SC-CR. Finally, previous analysis of *rec-8* animals showed that the SC-CR assembles between the two sisters, each forming an axis (Cahoon *et al*, 2019). The SC-CR in *rec-8* animals, like in wildtype animals, is therefore also between the DNA molecules undergoing an exchange.

The elevated sister exchanges in three conditions with different synaptonemal complex conformations *– syp*□*1*^*K42E*^, *ieDf2* and *rec*□*8 –* suggests that the SC-CR does not rely on a specific positioning of DNA molecules relative to one another to promote exchanges.

### Recombination nodules form throughout the SC-CR

As an orthogonal way to assess a potential role for the position of the SC-CR relative to the exchanged DNA molecules, we localized repair intermediates by assessing the position of recombination nodules. While previous analysis of hypotonically-treated samples suggested that recombination nodules adopt a stereotypical organization in the middle of the SC-CR (Woglar & Villeneuve, 2018), we wanted to localize recombination nodules in 3-dimensionally preserved samples.

We visualized COSA-1-GFP relative to the axes using STED microscopy. In wild-type animals, where crossovers form at recombination nodules, some foci localized between the axes in the middle of the SC-CR, but many were on one of the axes or off-center (Fig. 6). Foci were approximately Gaussian-distributed, with a full width at half maximum (FWHM) of 58% the inter-axis distance (95% CI of 50-68%; N = 94). This variation was approximately four-fold greater than the localization precision (Fig. 6B and S2). A similar, though more broadly distributed, localization pattern was observed for *ieDf2* animals, where sister exchanges, rather than inter-homolog crossovers, are generated at recombination nodules. In this case, foci were distributed with a FWHM of 85% the inter-axis distance (95% CI of 73-102%; N = 72), even though the inter-axis distance was similar to that in wildtype animals (Fig. 5 and Methods). Notably, 18% of foci localized outside the axes (Fig 6A, bottom), significantly more than the 5% observed in wildtype animals (Fisher’s exact test, p = 0.025).

**Figure 6:**
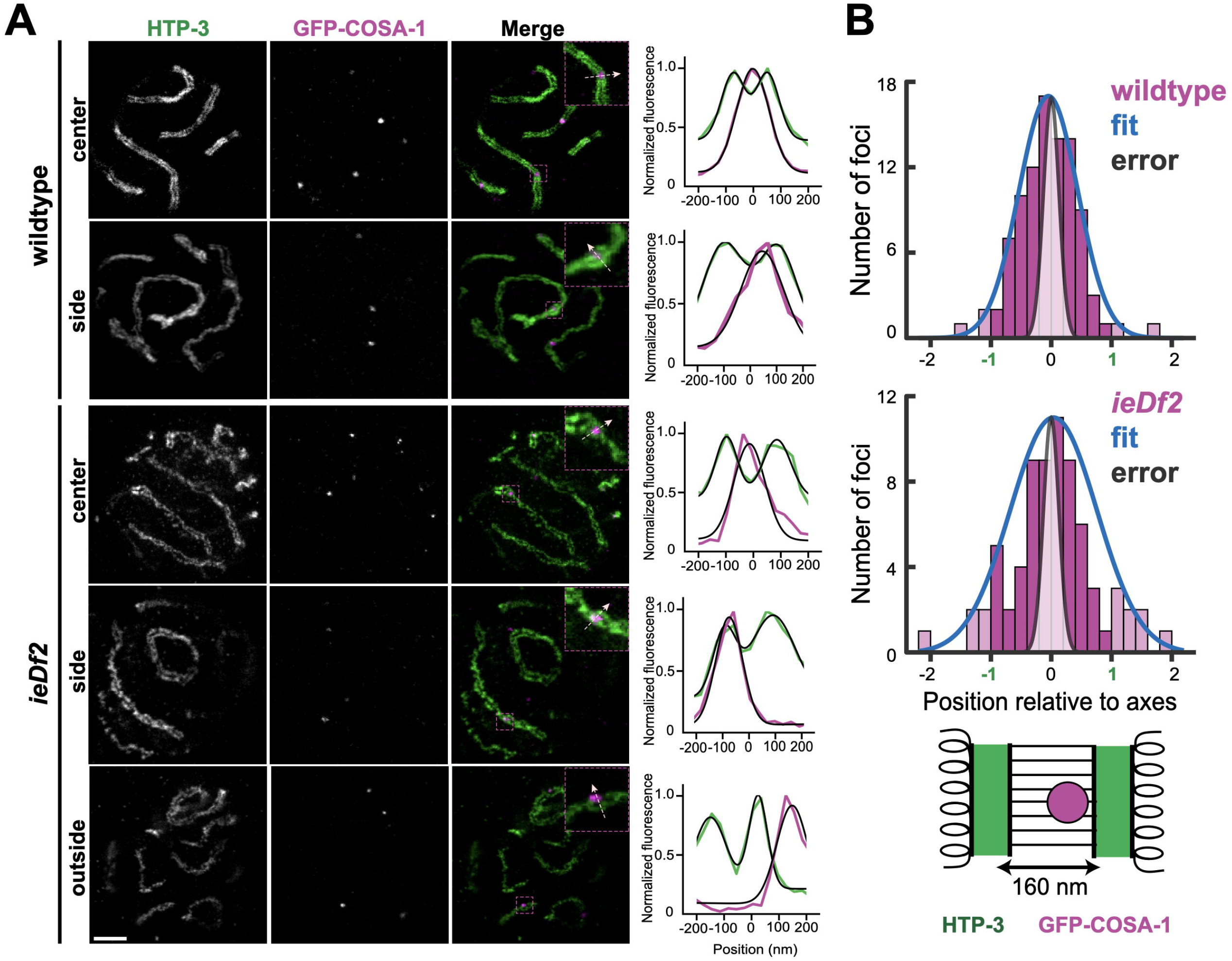
Recombination nodules do not form exclusively in the middle of the SC-CR. A. STED images of nuclei from the wildtype and *ieDf2* worms immunolabeled for GFP-COSA-1 (magenta) and HTP-3 (green). Right, intensity distributions extracted from line profiles of merged images (insets). Black lines denote fits to either one (GFP-COSA-1) or a sum of two (HTP-3) Gaussian distributions plus constant background. Scale bar = 1µm. B. Histograms of GFP-COSA-1 focus center positions for wildtype (n = 94) and *ieDf2* (n = 72). Values of 1 or -1 indicate localization on the meiotic axes, while a value of 0 indicates localization at the center of the SC-CR. Magnitudes >1 or <-1 (lighter bars) indicate GFP-COSA-1 center position outside the axes. Blue curves represent a Gaussian fit to the data, while the gray distribution represents estimated localization error (Fig. S2); for clarity, curves are displayed with the same maximum value as each histogram. Bottom, schematic diagram of synaptonemal complex geometry, with axes marked in green and the GFP-COSA-1 focus in magenta.

Our analysis indicates that the SC-CR directs the positioning of recombination nodules, and thereby the outcome of repair, independently of the position of the engaged DNA molecules. Rather than being in the middle of the SC-CR, the localization of recombination nodules is consistent with their formation throughout the volume of the SC-CR or at the top or bottom surface of the SC-CR.

## Discussion

Here we have documented correspondence between recombination nodules and sister exchanges in multiple independent scenarios (Figs. 1-2), dependency of most sister exchanges on recombination nodules (Fig. 3), and regulatory interplay between crossovers and sister exchanges (Fig.4). These data indicate that although only crossovers are known to form at recombination nodules, the same molecular machinery also regulates sister exchanges. An important implication of this finding is that the execution of the exchange/non-exchange decision likely relies on common features of inter-homolog and inter-sister events – e.g., repair intermediates like double Holliday junctions – rather than on their distinctive organization within meiotic chromosomes. While inter-homolog-specific processing suggests that recombination nodules will be positions strictly between the two homologs (Woglar & Villeneuve, 2018), we find that recombination nodules do not localize at a stereotypical location relative to the synaptonemal complex or to the two DNA molecules undergoing an exchange (Figs. 5 and 6). This supports a proximity-based mechanism for recombination nodule regulation by the synaptonemal complex – a lingering mystery due to the multiple, inter-dependent meiotic roles of the synaptonemal complex.

### Regulation of crossovers and sister exchanges

Our work establishes a two-tiered mechanism regulating homologous recombination during *C. elegans* meiosis. First, homolog bias directs most repair events to engage the homolog as a template. It is only in the absence of the homolog that the repair machinery resorts to using the sister chromatid as a template (MacQueen *et al*, 2005). The second layer independently carries out the exchange/non-exchange decision: while the vast majority of events form non-exchanges, those repair events designated by recombination nodules avoid this default fate and form exchanges. Together, these two tiers tightly regulate crossovers to ensure accurate chromosome segregation while channeling all other sister- and homolog-directed events into a non-exchange fate.

An open question is whether limiting the number of meiotic sister exchanges has a functional importance. Non-exchanges between sisters may be preferred due to the structural similarity between inter-sister and inter-homolog repair intermediates. The tight regulation on the number of crossovers may, as a byproduct, channel inter-sister events into the default, non-exchange fate. A non-mutually exclusive idea is that sister exchanges are deleterious, especially when occurring in regions rich in sequence repeats, and actively avoiding them helps protect genome integrity in the germline.

Our data indicates many commonalities in SC-CR-mediated regulation of crossovers and sister exchanges – among them correspondence with the number of recombination nodules, localization relative to the axes, and co-regulation *in cis*. However, it also points to potential differences. These include the localization of a minority of inter-sister recombination nodules outside the SC-CR (Fig. 6), and the seemingly lower efficiency of forming recombination nodules in *ieDf2* animals, where only four out of the 12 SC-CR stretches in the nucleus harbor a recombination nodule (Fig. 2). In addition, minor differences in exchange efficiency between crossovers and sister exchanges may be revealed by future work to directly measure the exact number of DNA breaks in worms (Yu *et al*, 2016) – currently a matter of debate since the available proxies, such as RAD-51 foci, are indirect and conflate break number with repair dynamics. While we have mitigated this limitation by comparing scenarios where meiotic checkpoints are already activated, this gap prevented us from comparing exchange frequency per sister-directed repair event in the different scenarios we have analyzed.

### Regulation of recombination nodules by the SC-CR

Our data shows that the SC-CR plays a crucial role in recruiting ZMM proteins and promoting their coalescence into functional recombination nodules. Some ZMM factors have an intrinsic affinity for the SC-CR: the ZMM proteins ZHP-1-4 and Vilya in worms and flies, respectively, localize along the SC-CR before forming recombination nodules. ZMM factors also localize to chromatin-free assemblies of SC-CR material (called polycomplexes) in worms, flies and yeast (Shinohara *et al*, 2015; Rog *et al*, 2017; Zhang *et al*, 2018; Lake *et al*, 2015; Tsubouchi *et al*, 2006; Voelkel-Meiman *et al*, 2019). Our work shows that recruitment of ZMM proteins to the SC-CR, and their coalescence into recombination nodules, promotes the formation of exchanges independently of the role of the SC-CR in bringing homologs together. We propose that SC-CR-mediated recruitment acts in concert with the affinity of ZMM members, like the MutS□complex Msh4-5 or the Zip2-Zip4-Spo16 complex, for specific repair intermediates (De Muyt *et al*, 2018; Snowden *et al*, 2004). While the molecular details of SC-CR–ZMM interactions in worms are unknown, recent work in budding yeast identified an interaction interface between the ZMM protein Zip4 and the SC-CR component Ecm11 (Pyatnitskaya *et al*, 2022).

Surprisingly, we find that the conserved conformation of the synaptonemal complex is not necessary for its ability to promote formation of recombination nodules (Fig. 5). Rather than being located in the middle of the SC-CR, recombination nodule distribution were distributed throughout its width (Fig. 6). This suggests that the SC-CR acts to concentrate ZMM proteins, and later to create a local environment in its vicinity that is conducive to coalescence of ZMM proteins into recombination nodules. Consistent with this idea is the location of dense recombination nodules outside the SC-CR in electron-micrographs of meiotic chromosomes in flies (Carpenter, 1975); and the formation of foci of ZMM proteins at the periphery of polycomplexes, which resemble recombination nodules (Shinohara *et al*, 2015; Rog *et al*, 2017; Zhang *et al*, 2018; Voelkel-Meiman *et al*, 2019; Tsubouchi *et al*, 2006).

In addition to its proximity-based role in positioning recombination nodules, the SC-CR may play additional roles in regulating meiotic exchanges. In *ieDf2* worms, where the DNA molecules undergoing exchange were to one side of the SC-CR, a minority of recombination nodules localized outside the SC-CR (Fig. 6). This points to a potential role for the conformation of the SC-CR relative to chromatin in tight regulation of recombination nodule position. Similarly, the presence of more recombination nodules than sister exchanges in *syp-1*^*K42E*^ animals (Figs. 1-2 and (Almanzar *et al*, 2021)) hints at a potential role of SC-CR conformation relative to the axes in ensuring the fidelity of exchange designation by recombination nodules.

What kind of material properties of the SC-CR and ZMM proteins could support these complex dynamics – initial colocalization followed by formation of juxtaposed yet distinct entities? The discovery that the SC-CR is a phase-separated compartment provides a potential mechanism. ZMM proteins initially interact more loosely with the SC-CR, either diffusing within it or forming a film around it. Coalescence of dynamic ZMM proteins into a recombination nodule reflects formation of a compartment with biophysical properties distinct from the SC-CR (Rog *et al*, 2017; Zhang *et al*, 2018). The coalescence and location of the recombination nodule compartment may be regulated by wetting or differential miscibility with the SC-CR (Feric *et al*, 2016; Wan *et al*, 2018). Repair intermediates, for which certain ZMM proteins have an affinity (De Muyt *et al*, 2018; Snowden *et al*, 2004), may direct the position of these coalescing events; in turn, these repair intermediates may be repositioned by forces exerted by interactions between the two compartments (Shin *et al*, 2018). Future work probing the 3-dimensional organization and dynamics of recombination nodules will test these mechanisms.

## Methods

### Worm strains and growing conditions

*C. elegans* worms were generated and cultured using standard conditions and protocols (Brenner, 1974). Worms were grown at 20°C, except worms carrying the *syp-1*^*K42E*^ allele, which were maintained at 15°C, but were grown at 25°C from hatching prior to experimentation. Auxin treatment was conducted as in (Almanzar *et al*, 2021) and (Zhang *et al*, 2015). *ieDf2* is a deletion of *him-8, zim-1, zim-2* and *zim-3* (Harper *et al*, 2011). The *ieDf2, syp-1*, and *syp-2* alleles were maintained as balanced strains, with homozygous progeny picked prior to experimentation. *syp-1(me17)* balanced by nT1 was used for the *nT1/+* experiments.

### Sister exchange analysis

EdU treatment, click chemistry and sister exchange quantification were conducted as in (Almanzar *et al*, 2021, 2022). Total number of sister exchanges (Fig. 2B) was extrapolated from the exchange rates on paired and unpaired chromosomes based on 6 paired chromosomes in wildtype animals, 5 paired and 2 unpaired chromosomes in *him-8* and *zim-2* animals, and 12 unpaired chromosomes in *ieDf2, syp-1, syp-2* and *syp-1*^*K42E*^ animals.

### Confocal microscopy

Immunofluorescence was performed as previously described (Phillips *et al*, 2009; Gordon *et al*, 2021), using ProLong Glass Antifade Mountant. All confocal microscopy images (Figs. 1, 2, 3, 4 and S1) were taken using a Zeiss LSM 880 confocal microscope equipped with an AiryScan and using a x63 1.4 NA oil immersion objective. Images were processed using Zen Blue 3.0. Partial maximum intensity projections are shown throughout.

### STED microscopy

Gonads were stained as previously reported (Phillips *et al*, 2009; Gordon *et al*, 2021) with the following modifications: for fixative, we used final concentration of 2% formaldehyde diluted from 16% ampules opened immediately prior to dissection; in addition, DAPI staining was omitted and MOUNT LIQUID (Aberrior) was used as mounting media. STAR RED and AlexaFluor 594 were used for the two STED channels (see Antibodies, below). Images were acquired with an Aberrior STEDYCON mounted on a Nikon Eclipse Ti microscope with a 100x 1.45 NA oil immersion objective. Images shown are a single z-section.

### Antibodies

The following antibodies were used: guinea-pig anti-HTP-3 (1:500, (Hurlock *et al*, 2020)), rabbit anti-SYP-5 (1:500, (Hurlock *et al*, 2020)), rabbit anti-SYP-2 (1:500, (Hurlock *et al*, 2020)), rabbit anti-RAD-51 (1:10,000, (Harper *et al*, 2011)), mouse anti-GFP (1:2,000, Roche), mouse anti-FLAG (1:500, Sigma-Aldrich), Cy3 AffiniPure Donkey anti-guinea-pig (1:500, Jackson Immunoresearch), 488 AffiniPure Donkey anti-mouse (1:500, Jackson Immunoresearch), 488 AffiniPure Donkey anti-rabbit (1:500, Jackson Immunoresearch), 594 AffiniPure Donkey anti-guinea-pig (1:200, Jackson Immunoresearch), 594 AffiniPure Donkey anti-mouse (1:200, Jackson Immunoresearch), Goat anti-rabbit STAR RED (1:200, Abberior), Goat anti-mouse STAR RED (1:200, Abberior) and Donkey anti-guinea-pig STAR RED (1:200, Abberior).

### Quantification of RAD-51 and GFP-COSA-1 foci

The mid-pachytene region of gonads stained with anti-RAD-51 antibodies were imaged. Maximum intensity projections spanning whole nuclei were set to the same threshold values across all genotypes to remove background staining. Foci from nuclei in late pachytene were counted. At least 3 gonads in each genotype were counted. GFP-COSA-1 was quantified by staining with an anti-GFP antibody, and scoring late-pachytene foci, as previously reported (Yokoo *et al*, 2012; Almanzar *et al*, 2021).

### Quantification of inter-axis width

Line scans across chromosomes in frontal view, where the axes and SC-CR were in the xy-plane, were performed using ImageJ. Following background subtraction (using the pixel with the lowest signal along the line scan), fluorescent measurements were normalized so that the pixel with the highest fluorescence is set to 1. Fluorescent plots were manually aligned so that the inter-axis minimum (for wildtype and *ieDf2*) or maximum (for *syp*□*1*^*K42E*^) were at the 0 μm point.

### Quantification of GFP-COSA-1 foci position relative to the SC-CR

To estimate the position of recombination nodules, we drew line profiles of the fluorescence intensity of GFP-COSA-1 and HTP-3, using the positions of SYP-2, which localizes in the center of the SC-CR (Fig. S2; (Köhler *et al*, 2020)), as an upper bound of our localization precision.

Fitting of GFP-COSA-1 and SYP-2 position relative to HTP-3 axis locations was performed using custom scripts written in MATLAB (The Mathworks; scripts available upon request). Briefly, line profiles were generated from STED data using the line profile function of Fiji (Schindelin *et al*, 2012) with a three-pixel wide (∼75 nm) line average. Fits to each line profile were performed using the lsqnonlin function of MATLAB, minimizing the residuals between the line profile data *y*(*x*) and a model defined by 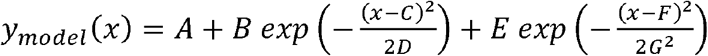 for free parameters A-G (or A-D for a single Gaussian). Initialization of the fit was performed by finding local maxima of *y*(*x*). To avoid overfitting of background for the double Gaussian, only data within 160 nm of either local maximum were fit. GFP-COSA-1 (or SYP-2) positions were normalized by subtracting the midpoint position of the two fit axis positions and dividing by the inter-axis distance of that profile; the sign of the distance was selected randomly for each measurement. Measured inter-axis distances were similar in GFP-COSA-1 experiments in wildtype and *ieDf2* worms, and in SYP-2 experiments, with mean values of 161 ± 25 nm (N = 94), 156 ± 27 nm (N = 72), and 162 ± 17 nm (N = 10), respectively (errors: standard deviation). Distributions of GFP-COSA-1 and SYP-2 positions within the inter-axis coordinate system were fit using maximum likelihood estimation to a Gaussian distribution, with full widths at half maximum of 0.58 (0.50,0.68), 0.85 (0.73,1.02), and 0.15 (0.11,0.29) relative to the distance between axes (ranges: 95% confidence interval).

### Statistical analysis

Chi-square and t-tests were conducted using a combination of the RStudio software package and GraphPad Prism.

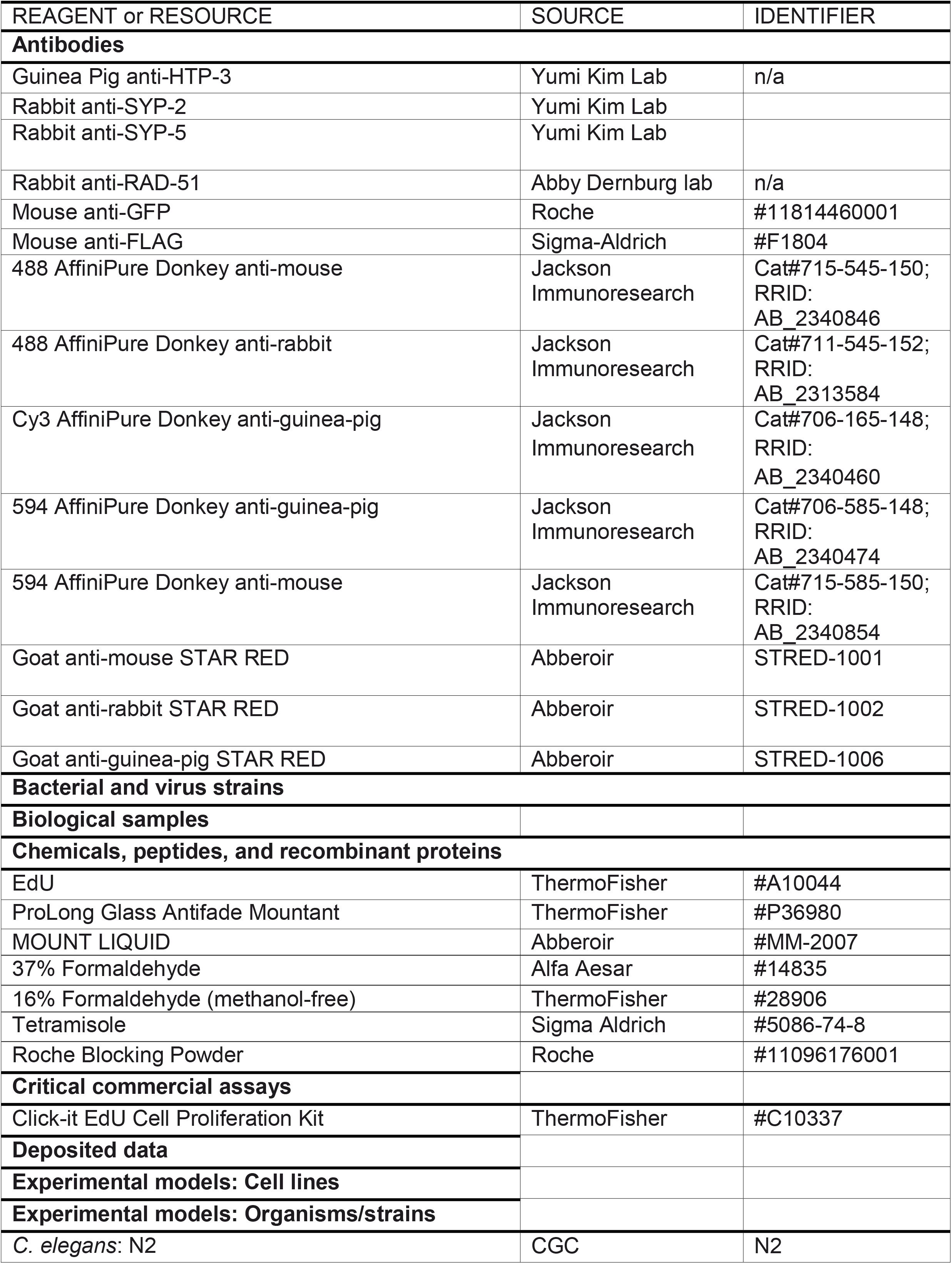
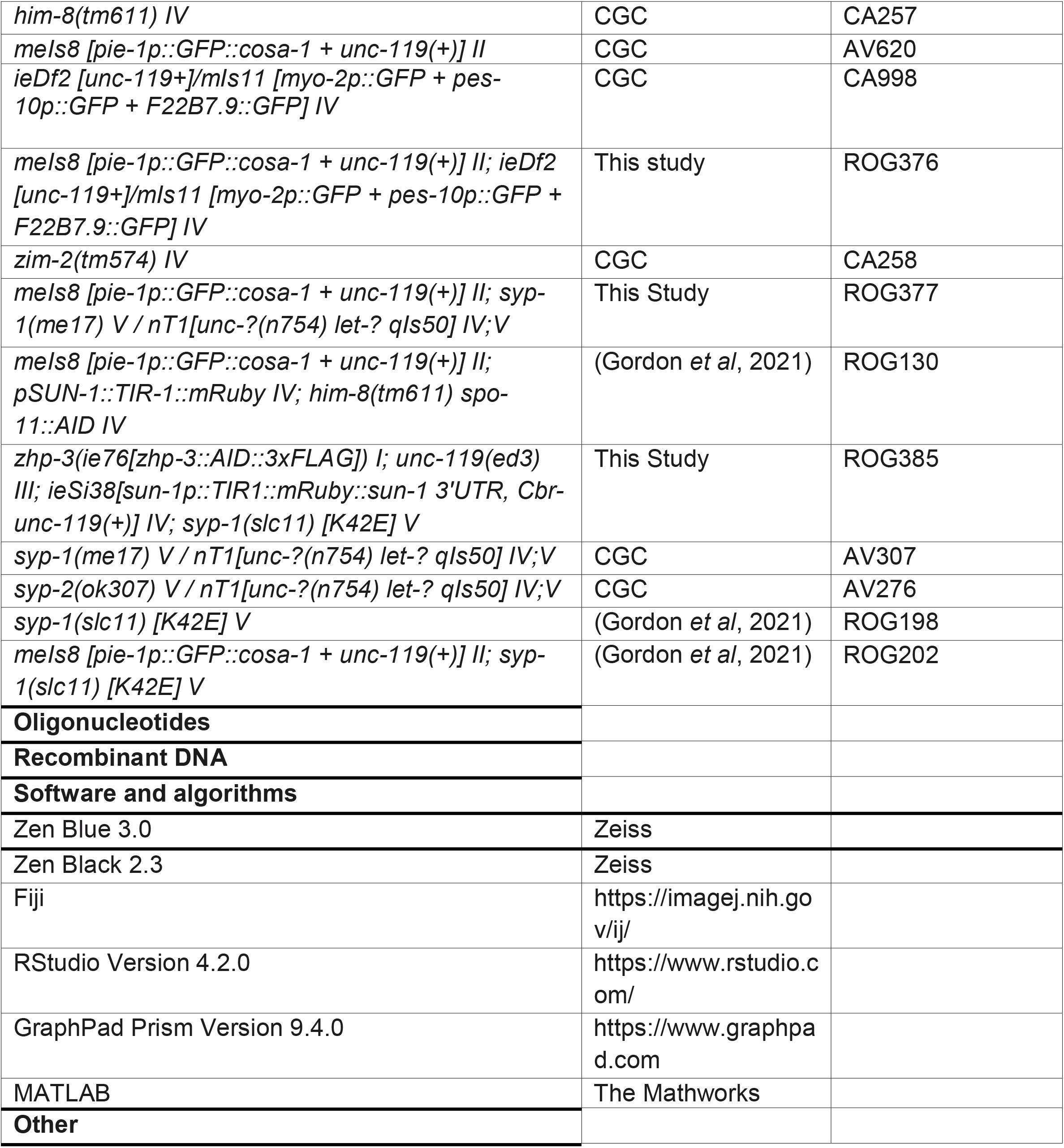

## Acknowledgements

We would like to thank members of the Rog Lab and Kent Golic for critical reading of this manuscript; Sara Nakielny for comments on the manuscript and editorial work; the University of Utah Cell Imaging facility for STED microscopy resources; and Yumi Kim and Abby Dernburg for antibodies. Worm strains were provided by the CGC, which is funded by NIH Office of Research Infrastructure Programs (P40 OD010440). LvD is The Mark Foundation for Cancer Research Fellow of the Damon Runyon Cancer Research Foundation (DRG-2372-19). This work was supported by a Genetics Training Grant T32GM007464 to DEA, and by a Pilot Project Award from the American Cancer Society, R35GM128804 grant from NIGMS, and start-up funds from the University of Utah to OR.

## Authors contributions

DEA performed most of the experimental work. SGG and CB performed the experimental work for Figures 5 and 6. DEA, SGG, CB and LvD analyzed the data. DEA and OR wrote the manuscript with input from all authors.

## Figure Legends

**Figure S1: RAD-51 foci are more numerous in *zim-2* compared with *him-8* or wildtype animals**

A. Representative images of mid-pachytene nuclei. Red, axis (anti-HTP-3 antibodies); yellow, RAD-51. The clusters of RAD-51 foci in *him-8* and *zim-2* nuclei are likely on the unpaired chromosomes (X and V, respectively; (MacQueen *et al*, 2005)). Scale bar = 1µm. B. Average number of RAD-51 foci per nucleus in mid-pachytene. *zim-2* exhibited significantly higher numbers of foci compared with wildtype and *him-8* animals (p<0.0001, Student’s t-test), consistent with a higher number of breaks.

**Figure S2: Localizing SYP-2 in the middle of the SC-CR to define localization error in STED experiments**

A. STED images of nuclei from the wildtype worms immunolabeled for SYP-2 (red) and HTP-3 (green). Right, intensity distributions extracted from line profiles of merged images (insets). Black lines denote fits to either one (SYP-2) or a sum of two (HTP-3) Gaussian distributions plus constant background. B. Histograms of SYP-2 focus center positions (n = 10). Values of 1 or -1 indicate localization on the meiotic axes, while a value of 0 indicates localization at the center of the SC-CR. Blue curve represent a Gaussian fit to the data. Bottom, schematic diagram of synaptonemal complex geometry, with axes marked in green and the antibodies targeting SYPL2 in red.

