## Supplementary figures and images for "Meiotic DNA exchanges are promoted by proximity to the synaptonemal complex"

### Fig S1

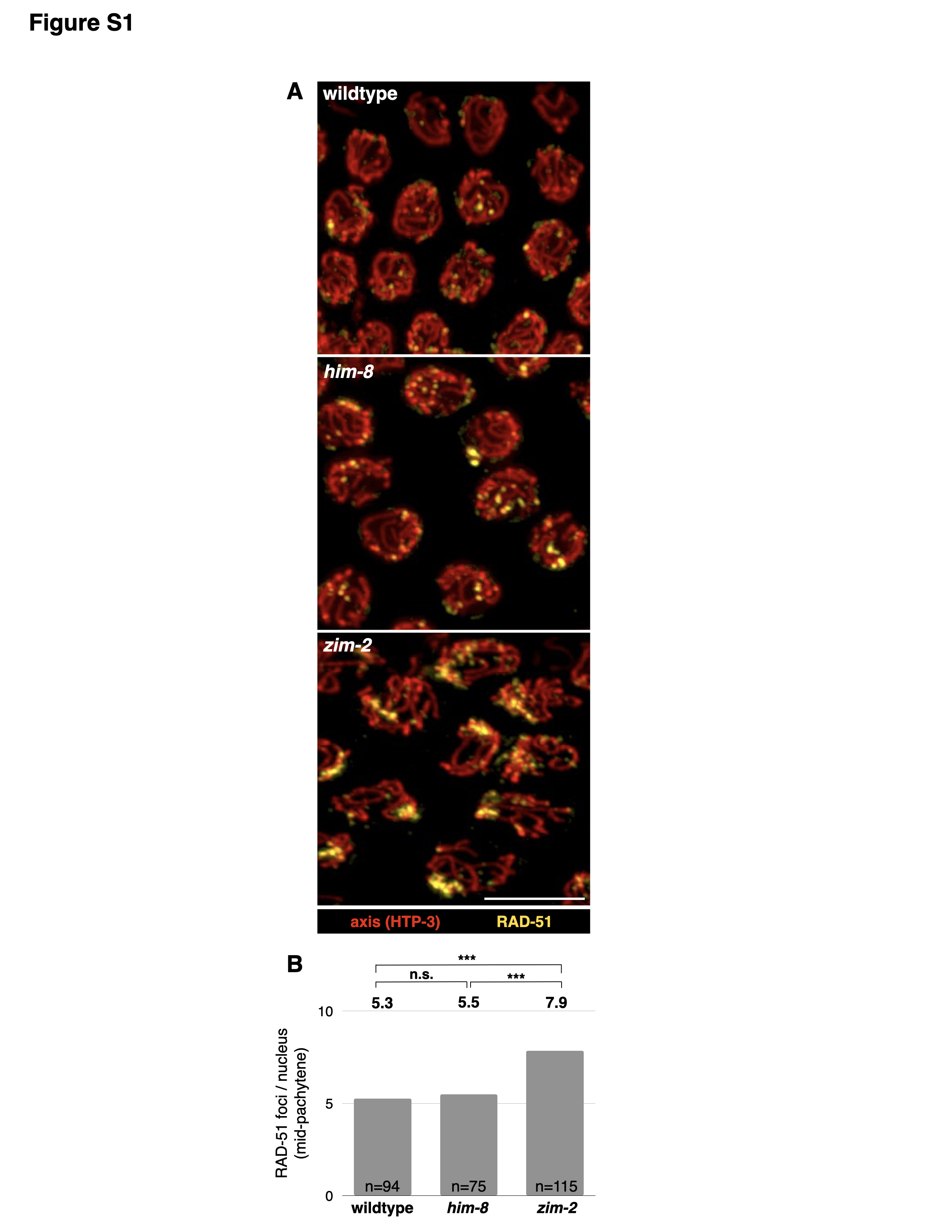

### Fig S2

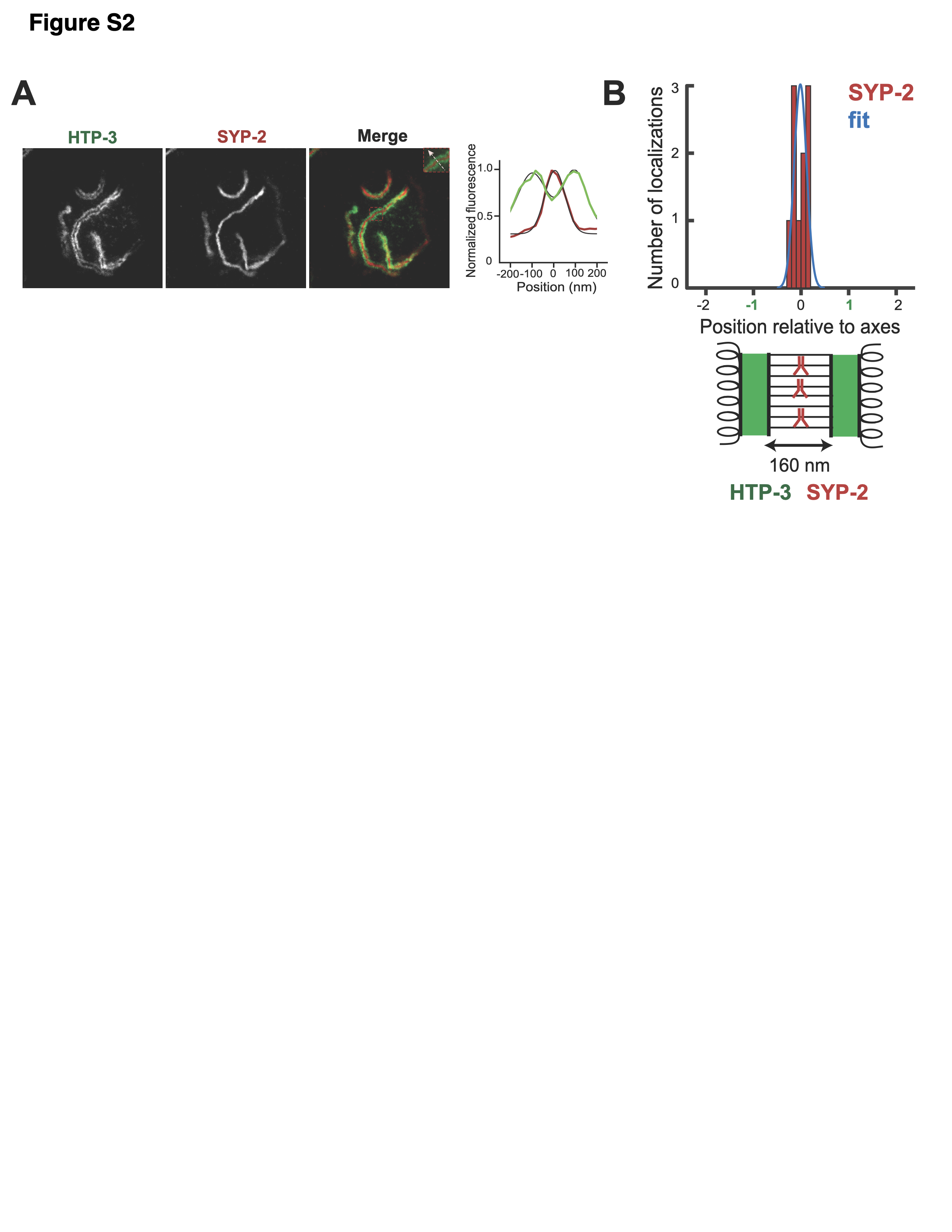
